# ASMC: investigating the amino acid diversity of enzyme active sites

**DOI:** 10.1101/2025.04.07.647545

**Authors:** Thomas Bailly, Eddy Elisée, David Vallenet

## Abstract

**Motivation:** The analysis of enzyme active sites is essential for understanding their activity in terms of catalyzed reaction and substrate specificity, providing insights for engineering to obtain targeted properties or modify the substrate scope. In 2010, a first version of the Active Site Modeling and Clustering (ASMC) workflow was published. ASMC predicts isofunctional clusters from enzyme families, based on structural modeling and clustering of active sites. Since then, structure- and sequence-based methods have developed considerably.

**Results:** We present here a redesign of the ASMC workflow. This new major version includes recent pocket prediction, structural alignment and clustering methods, as well as a refined amino acid distance matrix, thereby improving the relevance of results and reducing the need for laborious manual analysis to obtain relevant clusters. In addition, we have implemented multiple sequence alignment (MSA) as a possible input for the clustering step, along with an additional script to compare 2D and 3D active sites. Finally, the code has been unified from three to one programming language (Python) to facilitate its installation and maintenance. This new version of ASMC was evaluated on a set of protein families, resulting in overall better performances compared to its original version.

**Availability and implementation:** ASMC is supported on Linux operating system and freely available at https://github.com/labgem/ASMC, along with a complete documentation (wiki, tutorial).

## 1. Introduction

Identifying the residues involved in the reaction mechanism or substrate specificity is crucial for the study of enzyme families, as it helps define isofunctional subfamilies and provides insights for engineering enzymes to achieve targeted properties or modify their substrate scope. Over the years, various computational methods have been developed for the functionally relevant clustering of proteins, dealing with either hidden Markov models (SCI-PHY (Brown *et al*., 2007); GeMMA (Lee *et al*., 2009); FunFHMMer (Das *et al*., 2015)), phylogeny (AutoPhy (Ortiz-Velez 2024)), active site profiles (DASP (Fetrow, 2006); MISST (Harper *et al*., 2017)), genetic algorithms (de Lima *et al*., 2016) or 3D structures (TuLIP (Knutson *et al*., 2017)). In turn, the original Active Site Modeling and Clustering (ASMC) method (de Melo-Minardi *et al*., 2010) relies on homology modeling and active site classification to identify specificity determining positions (SDPs) and determine isofunctional groups. Over the years, ASMC has been employed by our team and collaborators to decipher the BKACE (Bastard *et al*., 2013), MetA/MetX (Bastard *et al*., 2017), CSL dioxygenase (Bastard *et al*., 2018) and AmDH (Mayol *et al*., 2019) protein families. We recently published a refined picture of the latter family (Elisée *et al*., 2024) in which we used a different reference structure (*Cfus*AmDH instead of AmDH4) for the ASMC analysis, which gave a better definition of the active site. Despite being available only upon request from the authors, ASMC has been cited approximately fifty times. Therefore, we believe that an open-source version would greatly benefit the scientific community. In addition, with advancements in the development of structure- and sequence-based methods, several components of the ASMC workflow could be improved. Furthermore, the hierarchical clustering method employed by ASMC (COBWEB (Fisher 1987) within WEKA software (Holmes, Donkin and Witten)) does not consider amino acid properties. This oversight can result in an excessive number of predicted subfamilies, necessitating manual investigation to ultimately obtain relevant clusters where active sites with chemically equivalent residues are grouped.

We present here a redesign of ASMC. In this new major version, we have updated the workflow by (i) unifying the code from three to one programming language (Python) to facilitate its installation and maintenance, (ii) implementing recent methods about pocket prediction, structural alignment and clustering with a refined amino acid distance matrix, (iii) making homology modeling with several reference structure or accepting external models provided by the user, (iv) using MSA input for the clustering, and other features detailed hereafter. We have evaluated ASMC on protein families mentioned above, resulting in overall better performances compared to its original version. Finally, ASMC is now freely available as an open-source software.

## 2. Methods and Implementation

ASMC is a precision tool to classify an enzyme family highlighting the diversity of amino acid residues making up the active site.

### 2.1 Architecture

The ASMC workflow has been improved for each of its four constituent steps described hereafter.

3D modeling: for each target sequence of the family, one reference structure is used as a template to build homology models through MODELLER (Webb and Sali, 2016) - if several reference structures are provided, the one with the best sequence identity is selected. This value should be at least 30% for a target sequence to be modeled (adjustable threshold). Prior to 3D modeling, a pairwise structural alignment between the target and its reference is generated using the SALIGN module (Madhusudhan *et al*., 2009). To reduce computation time, two models are built for each target and the one with the lowest DOPE score is selected. This step can be skipped if the user already has a set of 3D structures, for example taken from AI-based databases (Lin *et al*., 2023; Varadi *et al*., 2024), though care must be taken to ensure pocket accuracy for reliable active site identification and comparison.

Ligand-binding pocket search: based on carefully-chosen reference structure(s) (preferentially holo than apo form), the potential active site pocket is detected using P2RANK (Krivák and Hoksza, 2018). It uses machine learning to rapidly compute a ligandability score for pockets, defined from points distributed along the solvent-accessible protein surface. The user can then select one of the predicted pockets as reference (using P2RANK manually) or let the algorithm choose the one with the highest score.

Structural alignment: each homology model is aligned on its reference structure using US-align (Zhang *et al*., 2022). The resulting 3D sequence alignment is then used to retrieve the residue positions defined for the corresponding reference pocket. The same protocol is used for the 2D-based approach when an MSA is given as input.

Clustering: all active site sequences undergo an all-vs-all comparison to generate a score matrix, using a purpose-designed distance matrix for comparing active site residues (see Supplementary Note 1). The score matrix, scaled between 0 to 1, is then used by DBSCAN (Ester *et al*., 1996) to perform density-based spatial clustering guided by two parameters: *eps*, a distance measure to identify neighboring points, and *min_samples*, the minimum number of points for a region to be considered dense. The user can either let the algorithm define these values (see Supplementary Note 2) or play with their own.

### 2.2 Implementation

ASMC is Python-based (version ≥3.8), currently supported on Linux (tested on Fedora 40, CentOS 7, Ubuntu 24), and distributed on the GitHub platform as an open-source software. It requires four other easy-to-install bioinformatics tools (MODELLER, P2RANK, US-align) and some Python libraries: *plotnineseqsuite* (Cao *et al*., 2023), *biopython* (version ≥1.81), *numpy, scikit-learn, pyyaml, pillow*. Moreover, a Docker image is available to ease the installation.

The ASMC workflow can be launched in various ways depending on the user’s needs (Figure 1). Sequences and structures must be in FASTA and PDB format, respectively. Additional scripts are also available to aid in the analysis of the results.

**Figure 1.**
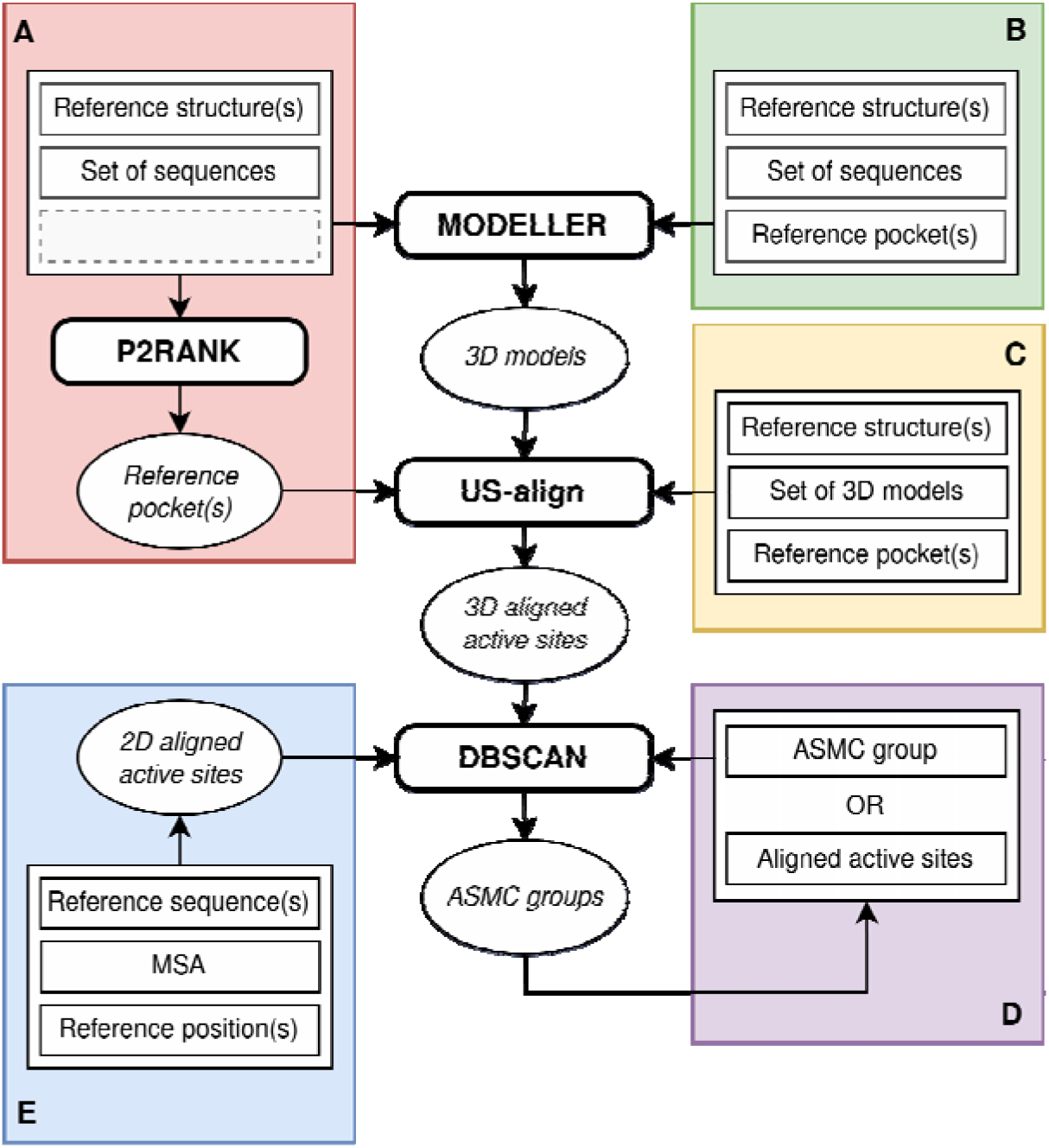
Schematic view of the updated ASMC workflow. Each colored frame represents a way to run ASMC: (**A**) whether the active site(s) is/are unknown or (**B**) known, (**C**) whether the 3D models are already available, (**D**) whether users need to re-cluster or sub-cluster a set of aligned active sites, or (**E**) whether the input is a multiple sequence alignment (2D-based approach). Software names are written in bold.

### 2.3 Benchmarking

Protein families previously analyzed using the original ASMC software were used to assess the relevance of the updated workflow’s results, in terms of cluster composition and sequence logos. These datasets comprise the BKACE (725 sequences), MetA (2,661 sequences), MetX (4,277 sequences) and AmDH (9,886 sequences) families.

## 3. Results

The updated version of ASMC automatically generates clusters consistent with those published and for which manual adjustments had been necessary (see Supplementary Figures S1-4, Supplementary Table S1 and Supplementary Note 3). This improved performance is primarily attributed to the distance matrix we developed, which enhances clustering of active site residues by more effectively accounting for their physicochemical similarities and differences.

ASMC was also able to extract active site sequences from the “non-active” groups of BKACE, AmDHs and MetX (Supplementary Figures S1-3), which is noteworthy given the presence of residues that are also involved in the active groups. Remarkably, ASMC succeeded in identifying the G3 group of the AmDH family (∼50 sequences among 9k+), which previously had to be manually curated. Finally, MetA clusters were sufficiently homogeneous to be separated by ASMC under specific residue positions (Supplementary Figure S4).

This new version of ASMC offers a concise overview of the amino acid diversity of active sites by greatly reducing the number of predicted clusters while providing, unlike its original version, a re-clustering option to further subdivide and explore groups of interest. This was performed, for example, for the largest groups in the BKACE and MetX families and revealed sub-clusters similar to those previously published, as well as new clusters with a refined amino acid profile (Supplementary Figures S5-6).

Experience has shown that ASMC clusters are more accurate when family entries share more than 30% of identity with reference structure protein sequences. Below this value, analysis should be carried out with caution and we recommend cross-checking with a MSA-based workflow to compare the results.

In terms of performance, the ASMC speed is limited by the homology modeling step, which logically depends on the number of target sequences. Reducing the number of models per target sequence from fifty to two (the default option value) had no impact on the overall results for the protein families studied, thus saving time.

## 4. Conclusions

ASMC has been updated to classify homologous enzyme sequences based on 2D or 3D active site residues, and to facilitate comparison of both approaches in order to evaluate the relevance of the active site profiles. Its use has been simplified and reduced to a single command line, offering numerous options for running ASMC according to the user’s needs. We believe this user-friendly version will be useful for quickly inspecting enzyme active sites and, for example, getting clues about residues that might be critical regarding substrate specificity, with the support of docking analysis.

## Funding

This project was supported by the French National Research Agency (ANR), through the ALADIN (ANR-21-ESRE-0021) project, the French Alternative Energies and Atomic Energy Commission (CEA), the National Centre for Scientific Research (CNRS) and the University of Evry Val d’Essonne - University of Paris-Saclay.

## Conflict of Interest

none declared.

## Data availability

The updated version of ASMC, described in this paper, is available under a CeCILL license from https://github.com/labgem/ASMC. Data used and generated in this application can be accessed through the Zenodo repository (https://doi.org/10.5281/zenodo.13483303). It contains the ASMC outputs, the starting sequence sets and the list of sequences discarded due to significant gaps, for the BKACE, MetA and MetX families. The AmDH 3D models were downloaded from Zenodo at https://doi.org/10.5281/zenodo.7889419.

## Notes

https://doi.org/10.5281/zenodo.13483303

### Competing Interest Statement

The authors have declared no competing interest.

