## Supplementary for "ASMC: investigating the amino acid diversity of enzyme active sites"

#### SUPPLEMENTARY TABLES

**Supplementary Table S1.** Number of clusters obtained with the original ASMC and its updated version. Published clusters correspond to manually-inferred clusters based on the hierarchical tree obtained with the ASMC original version. The columns ‘min’ indicates the minimum number of sequences required to form a cluster – for the updated ASMC, it corresponds to the ‘min_samples’ DBSCAN parameter (see Supplementary Note 2).

| | **Original ASMC (de Melo-Minardi *et al*. 2010)** | | | | **Updated ASMC (this article)** | |
|---|---|---|---|---|---|---|
| Protein families (size) | Total number of ASMC clusters* | min | Number of ASMC clusters > min | Published clusters | min | Total number of ASMC clusters** |
| BKACE (725) | 84 | 5 | 14 | 7 | 5 | 6 |
| MetA (2661) | 134 | 15 | 17 | 5 | 25 | 8 |
| MetX (4277) | 903 | 15 | 34 | 4 | 25 | 7 |
| AmDH (9886) | 1952 | 15 | 44 | 5 | 25 | 8 |

*direct ASMC output before application of the ‘min’ threshold.
**clusters larger than ‘min’ and obtained without any manual curation. Clusters smaller than ‘min’ are grouped in a miscellaneous cluster (annotated ‘-1’), which is included in the total count.

### SUPPLEMENTARY FIGURES

**Supplementary Figure S1.** ASMC clusters from the original ASMC and its updated version for the BKACE family. The number of sequences is given in brackets after the group name.

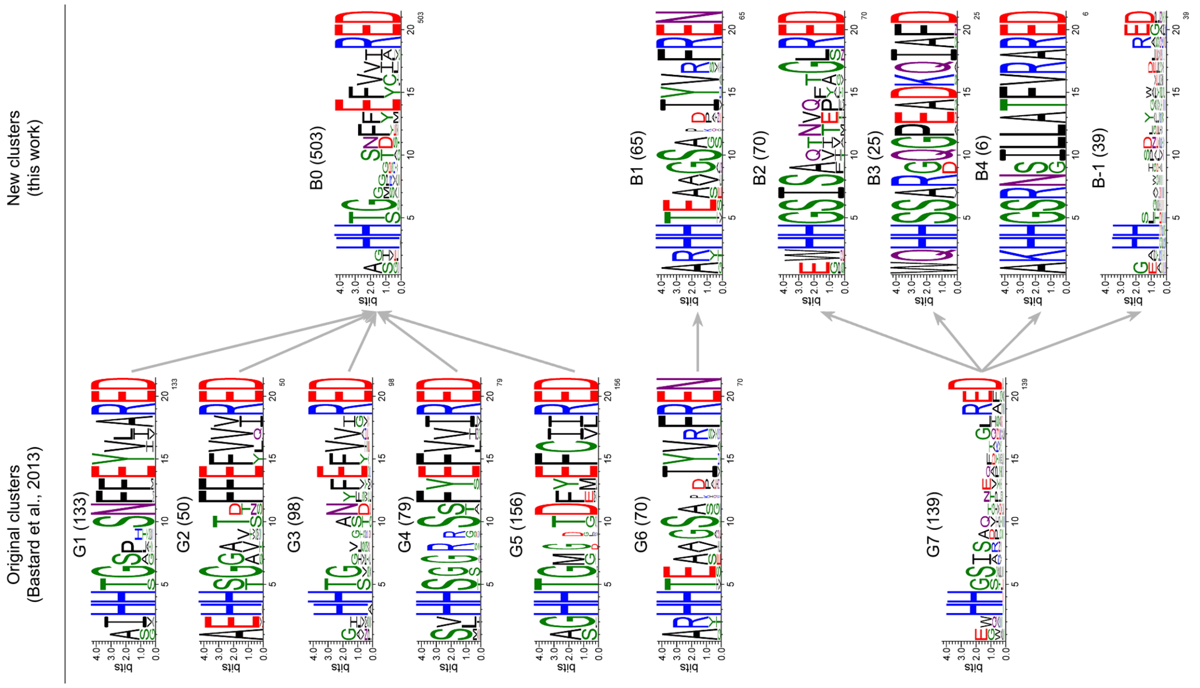

**Supplementary Figure S2.** ASMC clusters from the original ASMC and its updated version for the AmDH family. The number of sequences is given in brackets after the group name.

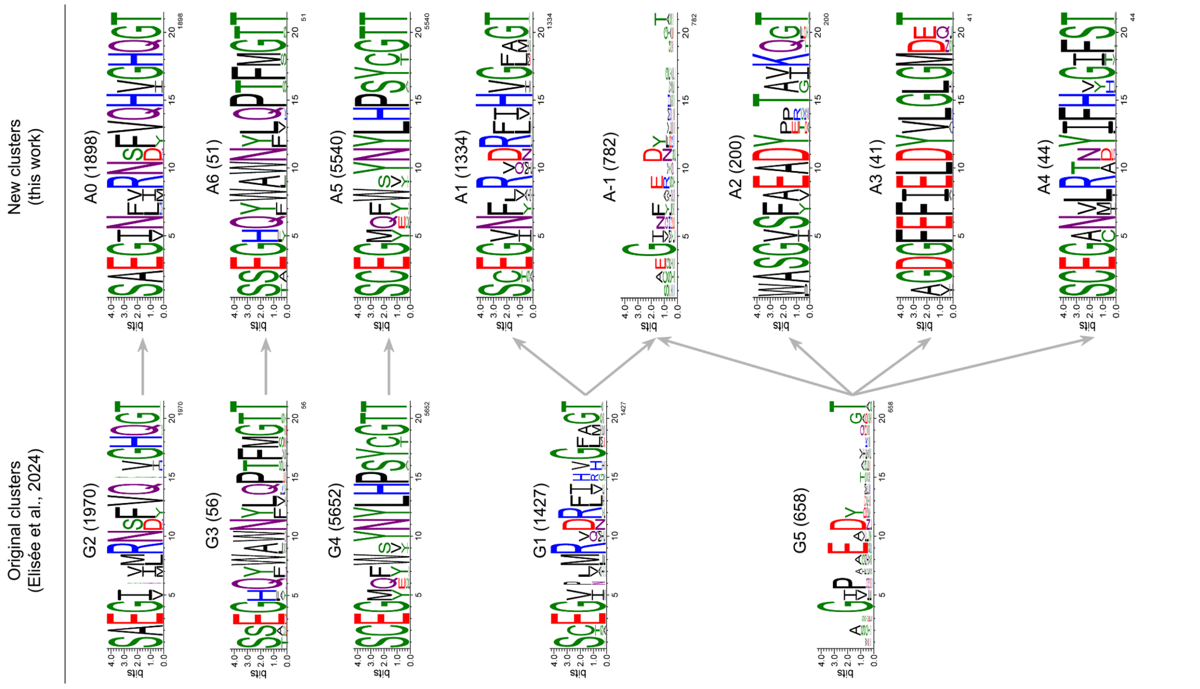

**Supplementary Figure S3.** ASMC clusters from the original ASMC and its updated version for the MetX family. The number of sequences is given in brackets after the group name.

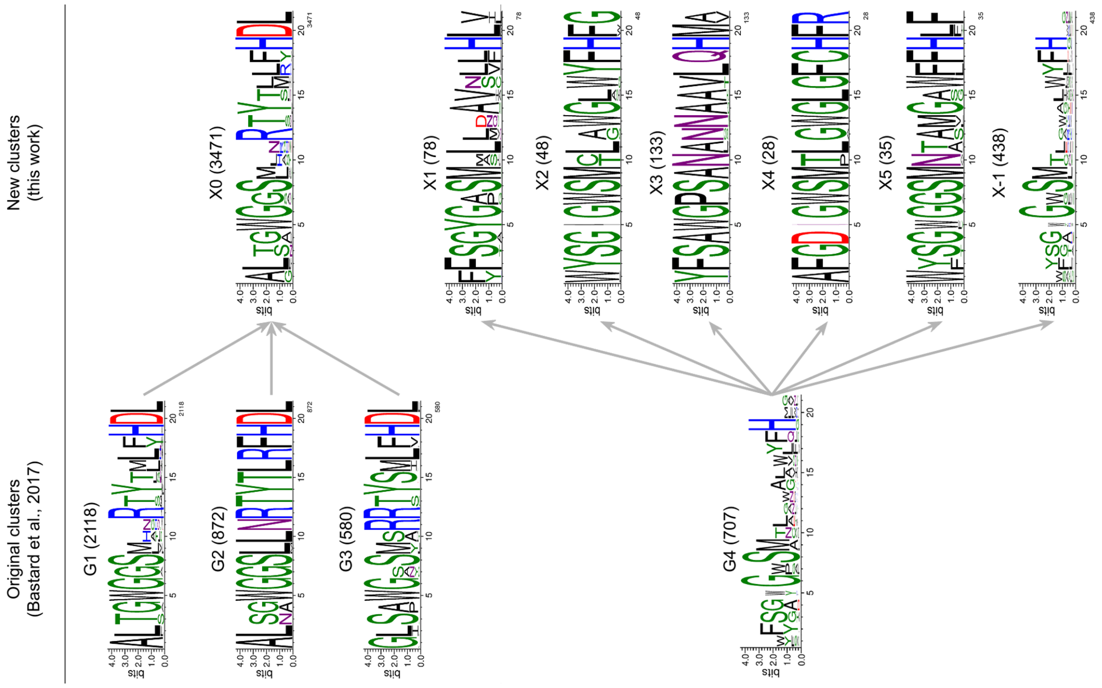

**Supplementary Figure S4.** ASMC clusters from the original ASMC and its updated version for the MetA family. The number of sequences is given in brackets after the group name.

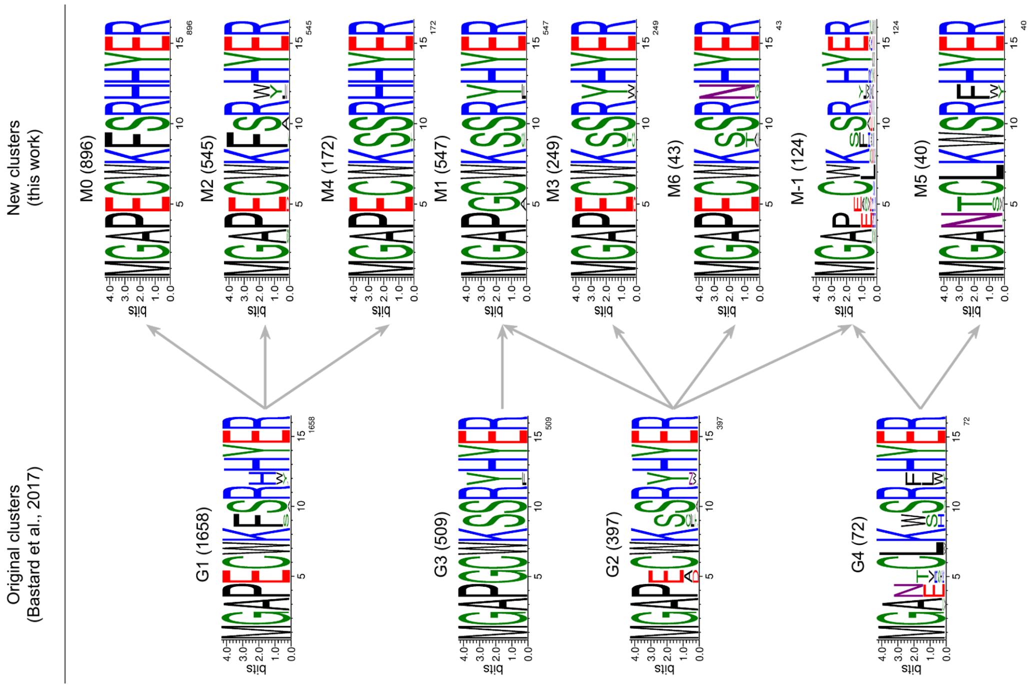

**Supplementary Figure S5.** Re-clustering of the B0 group (BKACE family). The number of sequences is given in brackets after the group name.

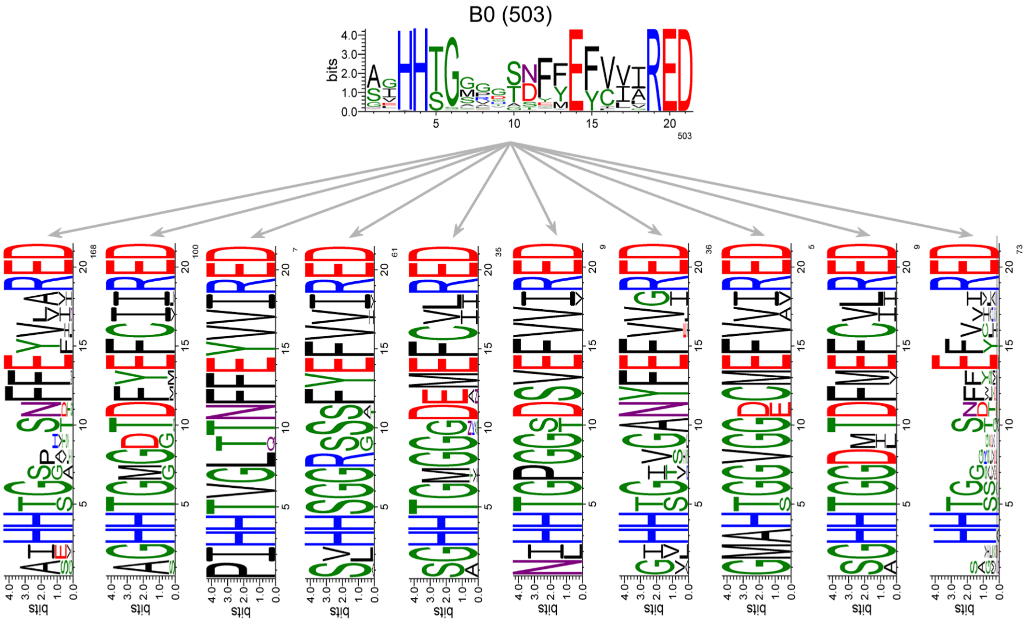

**Supplementary Figure S6.** Re-clustering of the X0 group (MetX family). The number of sequences is given in brackets after the group name.

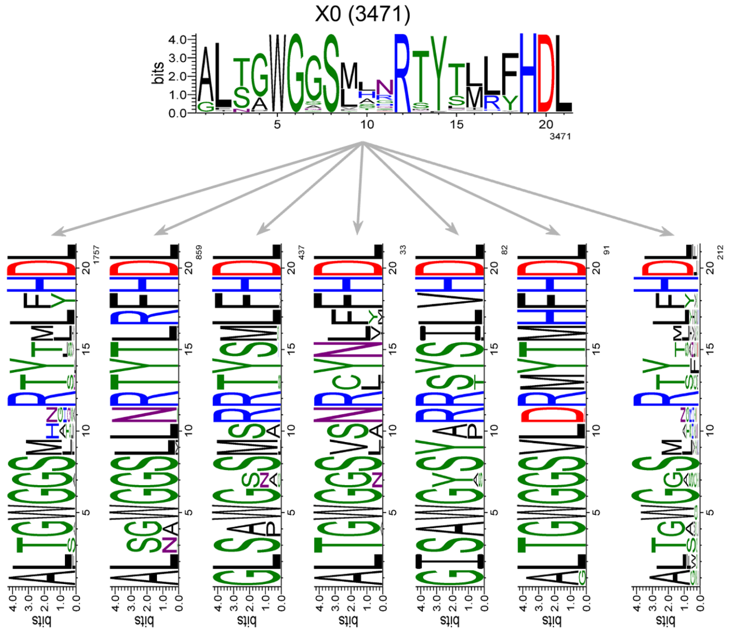

#### SUPPLEMENTARY NOTE 1

Generate the new distance matrix for classifying active site residues

We aimed to develop a distance matrix that accounts for the amino acid properties relevant to the analysis of the enzyme active sites. The Taylor classification (Taylor 1986) of amino acids generates overlapping groups, which is not practical in our case.

To achieve our goal, we used the AAindex database (Kawashima *et al.* 2007), which catalogs more than 500 physicochemical and biochemical properties of individual amino acids and pairs. We also reviewed the work of Then et al. (Then *et al.* 2020), which introduced a novel method for deriving an optimal classification of amino acids. Rather than eliminating features based on correlation thresholds, we chose to refine the 83 features identified by Then et al. (Then *et al.* 2020) (as detailed in Table S1 of their additional information) in a stepwise manner. This approach involved retaining only 7 features from the original 83 by removing those deemed irrelevant to the active site environment and/or those already correlated with more pertinent features. The correlation matrix for the 83 features is presented hereafter.

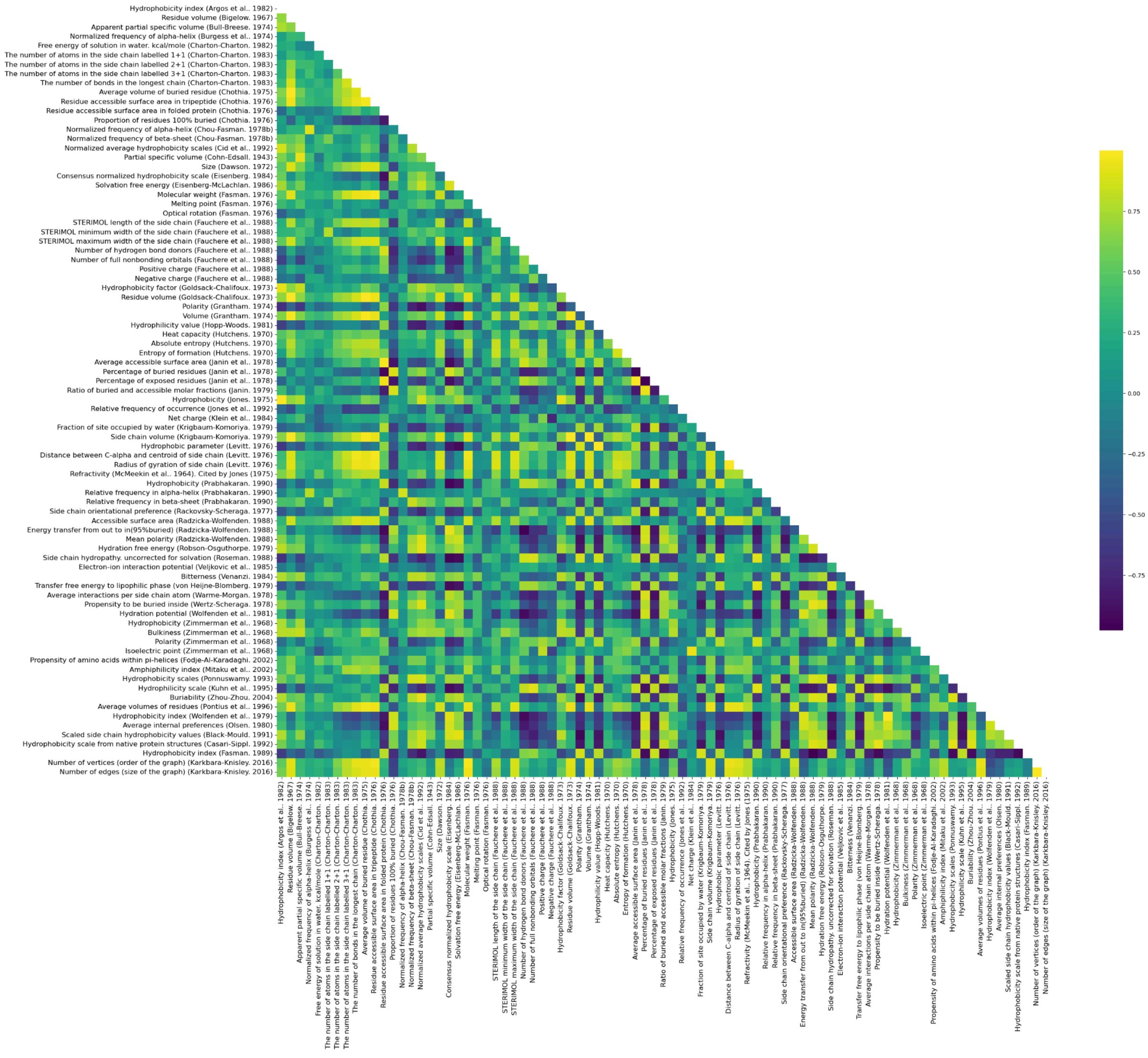

The AAindex values for the 7 selected features are shown in the following table.

| Aminoacids (1-letter code) | Free energy of solution in water. kcal/mole (Charton-Charton. 1982) | Number of hydrogen bond donors (Fauchere *et al*. 1988) | Net charge (Klein *et al*. 1984) | Side chain volume (Krigbaum-Komoriya. 1979) | Electron-ion interaction potential (Veljkovic *et al*. 1985) | Hydrophobicity scale from native protein structures (Casari-Sippl. 1992) | Aromatic |
|---|---|---|---|---|---|---|---|
| **A** | -0.368 | 0.0 | 0.0 | 27.5 | 0.03731 | 0.2 | 0 |
| **R** | -1.03 | 4.0 | 1.0 | 105.0 | 0.09593 | -0.7 | 0 |
| **N** | 0.0 | 2.0 | 0.0 | 58.7 | 0.00359 | -0.5 | 0 |
| **D** | 2.06 | 1.0 | -1.0 | 40.0 | 0.1263 | -1.4 | 0 |
| **C** | 4.53 | 0.0 | 0.0 | 44.6 | 0.08292 | 1.9 | 0 |
| **Q** | 0.731 | 2.0 | 0.0 | 80.7 | 0.07606 | -1.1 | 0 |
| **E** | 1.77 | 1.0 | -1.0 | 62.0 | 0.0058 | -1.3 | 0 |
| **G** | -0.525 | 0.0 | 0.0 | 0.0 | 0.00499 | -0.1 | 0 |
| **H** | 0.0 | 1.0 | 1.0 | 79.0 | 0.02415 | 0.4 | 1 |
| **I** | 0.791 | 0.0 | 0.0 | 93.5 | 0.0 | 1.4 | 0 |
| **L** | 1.07 | 0.0 | 0.0 | 93.5 | 0.0 | 0.5 | 0 |
| **K** | 0.0 | 2.0 | 1.0 | 100.0 | 0.0371 | -1.6 | 0 |
| **M** | 0.656 | 0.0 | 0.0 | 94.1 | 0.08226 | 0.5 | 0 |
| **F** | 1.06 | 0.0 | 0.0 | 115.5 | 0.0946 | 1.0 | 1 |
| **P** | -2.24 | 0.0 | 0.0 | 41.9 | 0.01979 | -1.0 | 0 |
| **S** | -0.524 | 1.0 | 0.0 | 29.3 | 0.08292 | -0.7 | 0 |
| **T** | 0.0 | 1.0 | 0.0 | 51.3 | 0.09408 | -0.4 | 0 |
| **W** | 1.6 | 1.0 | 0.0 | 145.5 | 0.05481 | 1.6 | 1 |
| **Y** | 4.91 | 1.0 | 0.0 | 117.3 | 0.05159 | 0.5 | 1 |
| **V** | 0.401 | 0.0 | 0.0 | 71.5 | 0.00569 | 0.7 | 0 |

Firstly, each of these 7 features was scaled independently between 0 and 1 since they have different units. We then calculated the cityblock distance using the SciPy Python package and several combinations of weights (default value = 1) were subsequently tested. The aminoacid residues were projected in 2D based on the distance matrix, using the MDS function in the scikit-learn Python package. Finally, a suitable 2D distribution was obtained for the physico-chemically similar amino acid residues, with the weights shown below.

| Features | Weight |
|---|---|
| Free energy of solution in water. kcal/mole (Charton-Charton. 1982) | 0.75 |
| Number of hydrogen bond donors (Fauchere *et al*. 1988) | 1 |
| Net charge (Klein *et al*. 1984) | 2 |
| Side chain volume (Krigbaum-Komoriya. 1979) | 1 |
| Electron-ion interaction potential (Veljkovic *et al*. 1985) | 1.5 |
| Hydrophobicity scale from native protein structures (Casari-Sippl. 1992) | 1 |
| Aromatic | 1.75* |

*This value is set at 1 for the histidine residue to favor its net charge rather than its aromaticity and to discriminate it from the other aromatic residues (Y, F, W).

The final distance matrix is presented below and is available on the GitHub repository (https://github.com/labgem/ASMC/blob/main/resources/AA_distances.tsv).

| | A | R | N | D | C | Q | E | G | H | I | L | K | M | F | P | S | T | W | Y | V |
|---|---|---|---|---|---|---|---|---|---|---|---|---|---|---|---|---|---|---|---|---|
| **A** | 0 | 14 | 5 | 11 | 6 | 7 | 10 | 2 | 10 | 5 | 4 | 10 | 4 | 10 | 3 | 4 | 5 | 12 | 11 | 3 |
| **R** | 14 | 0 | 11 | 16 | 15 | 9 | 18 | 15 | 10 | 15 | 14 | 6 | 11 | 15 | 14 | 10 | 9 | 18 | 17 | 14 |
| **N** | 5 | 11 | 0 | 12 | 10 | 4 | 7 | 4 | 10 | 5 | 5 | 8 | 7 | 13 | 5 | 5 | 5 | 13 | 12 | 4 |
| **D** | 11 | 16 | 12 | 0 | 11 | 9 | 5 | 13 | 18 | 15 | 14 | 15 | 11 | 16 | 11 | 8 | 7 | 17 | 17 | 13 |
| **C** | 6 | 15 | 10 | 11 | 0 | 8 | 13 | 9 | 14 | 7 | 8 | 15 | 5 | 9 | 8 | 6 | 6 | 11 | 10 | 7 |
| **Q** | 7 | 9 | 4 | 9 | 8 | 0 | 9 | 9 | 11 | 8 | 7 | 7 | 4 | 10 | 7 | 4 | 4 | 11 | 11 | 7 |
| **E** | 10 | 18 | 7 | 5 | 13 | 9 | 0 | 9 | 14 | 9 | 8 | 12 | 11 | 17 | 8 | 10 | 9 | 16 | 15 | 8 |
| **G** | 2 | 15 | 4 | 13 | 9 | 9 | 9 | 0 | 11 | 5 | 4 | 12 | 7 | 13 | 3 | 5 | 6 | 14 | 14 | 3 |
| **H** | 10 | 10 | 10 | 18 | 14 | 11 | 14 | 11 | 0 | 10 | 9 | 7 | 11 | 10 | 11 | 11 | 11 | 9 | 8 | 9 |
| **I** | 5 | 15 | 5 | 15 | 7 | 8 | 9 | 5 | 10 | 0 | 1 | 11 | 4 | 9 | 6 | 9 | 8 | 9 | 11 | 2 |
| **L** | 4 | 14 | 5 | 14 | 8 | 7 | 8 | 4 | 9 | 1 | 0 | 10 | 3 | 9 | 5 | 8 | 7 | 10 | 10 | 1 |
| **K** | 10 | 6 | 8 | 15 | 15 | 7 | 12 | 12 | 7 | 11 | 10 | 0 | 10 | 16 | 10 | 10 | 10 | 15 | 15 | 10 |
| **M** | 4 | 11 | 7 | 11 | 5 | 4 | 11 | 7 | 11 | 4 | 3 | 10 | 0 | 6 | 7 | 5 | 4 | 9 | 9 | 4 |
| **F** | 10 | 15 | 13 | 16 | 9 | 10 | 17 | 13 | 10 | 9 | 9 | 16 | 6 | 0 | 13 | 11 | 9 | 4 | 5 | 10 |
| **P** | 3 | 14 | 5 | 11 | 8 | 7 | 8 | 3 | 11 | 6 | 5 | 10 | 7 | 13 | 0 | 5 | 6 | 14 | 13 | 4 |
| **S** | 4 | 10 | 5 | 8 | 6 | 4 | 10 | 5 | 11 | 9 | 8 | 10 | 5 | 11 | 5 | 0 | 2 | 12 | 12 | 7 |
| **T** | 5 | 9 | 5 | 7 | 6 | 4 | 9 | 6 | 11 | 8 | 7 | 10 | 4 | 9 | 6 | 2 | 0 | 11 | 11 | 6 |
| **W** | 12 | 18 | 13 | 17 | 11 | 11 | 16 | 14 | 9 | 9 | 10 | 15 | 9 | 4 | 14 | 12 | 11 | 0 | 4 | 11 |
| **Y** | 11 | 17 | 12 | 17 | 10 | 11 | 15 | 14 | 8 | 11 | 10 | 15 | 9 | 5 | 13 | 12 | 11 | 4 | 0 | 11 |
| **V** | 3 | 14 | 4 | 13 | 7 | 7 | 8 | 3 | 9 | 2 | 1 | 10 | 4 | 10 | 4 | 7 | 6 | 11 | 11 | 0 |

#### SUPPLEMENTARY NOTE 2

How DBSCAN parameters are computed when the “auto” option is used

*min_samples*
The “auto” value is defined based on the number of input protein sequences. It will be equal to 5 or 25, if there are fewer or more than 1500 sequences, respectively.

*eps*
The value is derived from the distribution of the score matrix computed during the all-vs-all comparison step and supplied to DBSCAN. We considered the first quantile of this distribution and deducted 10% empirically to obtain the “auto” value.

#### SUPPLEMENTARY NOTE 3

Comparison of previously published ASMC groups with those obtained with the updated version of ASMC

*BKACE*

The group B0 gathers the previously published G1-5 groups and was relevant for a re-clustering step to evaluate if the updated ASMC was able to retrieve them. The group B1 is consistent with the group G6. Finally, the groups B-1, B3, B4 and B5 are consistent with the G7 group labeled as “not BKACE”. B3, B4 and B5 emerged from it due to amino acidic specific positions considered as sufficiently significant by the DBSCAN method to form new clusters.

*AmDHs*

The four main groups are retrieved without manual investigations (G1/A1, G2/A0, G3/A6, G4/A5). The group G5 is splitted into 4 groups, one outlier (A-1), one with an arginine residue (R) at position 9 like group G1 (A4), and two “not AmDH”-labeled groups (A2, A3) as there is no glutamate residue (E) at position 3, which is critical for catalysis.

*MetX*

The groups G1, G2 and G3 are merged into one group (X0) which is relevant for a re-clustering step, like B0 of the BKACE family. The G4 group, displaying a heterogenous sequence logo, has been split into 5 consistent groups (X1-5) and the outliers (X-1).

*MetA*

The group G1 is retrieved into three groups (M0, M2, M4) based on positions 9 and 12, results which support the relevance of our score matrix specially designed on amino acid properties within the active site. Similarly, the group G2 is retrieved into four groups (M1, M3, M6, M-1) based on positions 5, 9 and 12. The group G3 is consistent with the group M1, with few inputs from the group G2. Finally, the group G4 is retrieved into one cluster (M5) and the outliers (M-1), based on position 4 and 12.(The classification of amino acid conservation 1986)
